# Rapid, efficient auxin-inducible protein degradation in *Candida* pathogens

**DOI:** 10.1101/2023.05.17.541235

**Authors:** Kedric L. Milholland, Justin B. Gregor, Smriti Hoda, Soledad Píriz-Antúnez, Encarnación Dueñas-Santero, Bao Gia Vu, Krishna P. Patel, W. Scott Moye-Rowley, Carlos R. Vázquez de Aldana, Jaime Correa-Bordes, Scott D. Briggs, Mark C. Hall

## Abstract

A variety of inducible protein degradation (IPD) systems have been developed as powerful tools for protein functional characterization. IPD systems provide a convenient mechanism for rapid inactivation of almost any target protein of interest. Auxin-inducible degradation (AID) is one of the most common IPD systems and has been established in diverse eukaryotic research model organisms. Thus far, IPD tools have not been developed for use in pathogenic fungal species. Here, we demonstrate that the original AID and the second generation AID2 systems work efficiently and rapidly in the human pathogenic yeasts *Candida albicans* and *Candida glabrata*. We developed a collection of plasmids that support AID system use in laboratory strains of these pathogens. These systems can induce >95% degradation of target proteins within minutes. In the case of AID2, maximal degradation was achieved at low nanomolar concentrations of the synthetic auxin analog 5-adamantyl-indole-3-acetic acid (5-Ad-IAA). Auxin-induced target degradation successfully phenocopied gene deletions in both species. The system should be readily adaptable to other fungal species and to clinical pathogen strains. Our results define the AID system as a powerful and convenient functional genomics tool for protein characterization in fungal pathogens.

## INTRODUCTION

Historically, functional characterization of proteins has largely depended either on their biochemical isolation and characterization or on gene inactivation through random mutation or targeted chromosome editing followed by phenotypic observation. Permanent gene inactivation can significantly alter cellular physiology though, including activating adaptation mechanisms or selecting for compensatory mutations. More recently, transcriptional repression and RNA interference technologies have provided useful alternatives, for example in studying essential gene products (Bellí et al., 1998; Clemens et al., 2000; Kamath et al., 2003; Mnaimneh et al., 2004; Roemer et al., 2003). Transcriptional repression and RNA silencing methods are often slow-acting, particularly for stable proteins that must be fully degraded before phenotypes appear. In these cases, the observed phenotypes may not be directly linked to functions of the protein of interest, but rather an indirect consequence of its persistent loss of function. Moreover, the permanent or slow-acting nature of these methods makes them less applicable for mechanistic studies of dynamic cellular processes. The use of specific chemical inhibitors is an ideal way to study protein function that circumvents many of the problems associated with molecular genetic methods for reducing protein function. Inhibitors can be fast-acting, can achieve near complete loss of function, are often reversible, and in many cases are highly specific. However, not all proteins have functions like enzymatic activity that can be readily inhibited by small molecules, and effective, highly specific inhibitors are not available for most proteins.

The advent of inducible protein degradation (IPD) technologies has provided a “best of both worlds” option for protein functional characterization. In IPD systems a target gene is engineered using molecular genetic tools so that the encoded protein is functional and expressed at natural level but its proteolytic degradation can be rapidly triggered by an external stimulus. These systems can be applied, in principle, to any target protein yet have the speed, specificity, and efficacy of chemical inhibitors. Many inducible degradation systems have been developed for use in a variety of model organisms (Natsume and Kanemaki, 2017; Prozzillo et al., 2020). One of the most popular is the auxin-inducible degradation (AID) system (**Figure 1A**), based on the natural mode of action of the plant hormone, auxin. Auxins act as molecular glues that promote physical association of auxin binding domain (ABD) proteins with the SCF^Tir1^ ubiquitin ligase (Dong et al., 2021; Salehin et al., 2015; Schreiber, 2021). This results in polyubiquitination of the ABD protein and its subsequent recognition and proteolysis via the 26S proteasome. In 2009 Kanemaki and colleagues demonstrated that fusion of a target gene to the coding sequence of an ABD and expression of a plant Tir1 F-box protein in *S. cerevisiae* or cultured human cells allowed rapid degradation of the fusion protein simply by addition of the natural auxin, 3-indoleacetic acid (IAA), to the culture medium (Nishimura et al., 2009).

**Figure 1.**
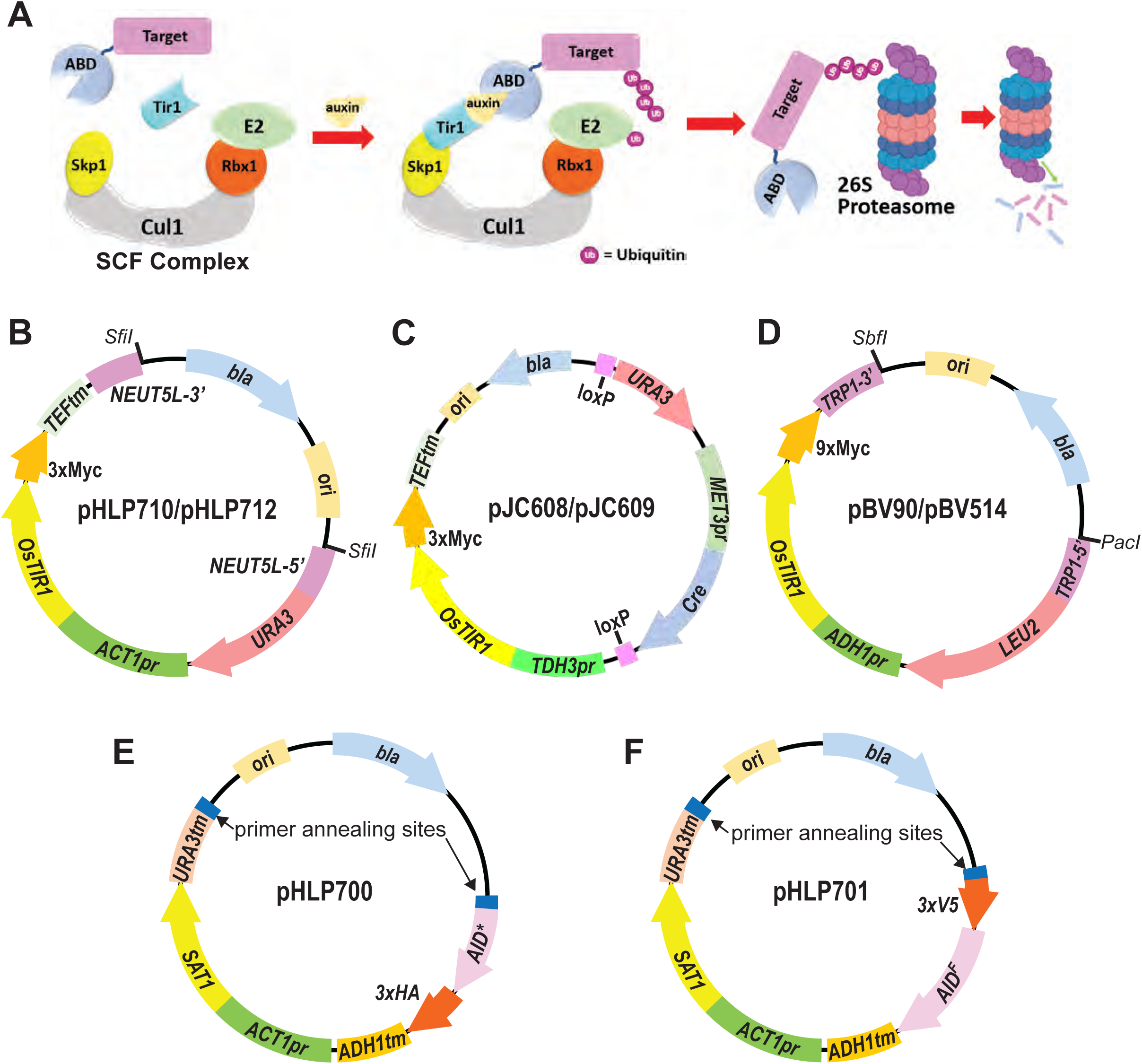
Overview of AID system and constructs for *Candida* strain engineering. **A)** AID system function requires 1) fusion of an auxin-binding domain (ABD), e.g., from *A. thaliana* IAA17, to a target protein and 2) ectopic expression of a plant *TIR1* gene, e.g., from *Oryza sativa*, that can interact with the host organism’s endogenous SCF E3 ubiquitin ligase complex. Addition of auxin to cells induces Tir1-ABD interaction and recruitment of the target protein to the SCF-E2 complex for polyubiquitination and subsequent degradation via the 26S proteasome. **B)** Plasmid containing *OsTIR1* integration cassette for *C. albicans*. Cassette bounded by *NEUT5L* homology regions can be excised by *SfiI* digest or amplified by PCR for transformation and integration at *NEUT5L* locus. **C)** Plasmid containing *OsTIR1* integration cassette with recyclable *URA3* marker. After integration at the desired locus, removal of methionine leads to Cre expression and recombination between loxP sites, excising *URA3* and Cre genes. **D)** Plasmid containing *OsTIR1* integration cassette for *C. glabrata.* Cassette bounded by *TRP1* homology regions can be excised by restriction digest or amplified by PCR for transformation and integration at *TRP1*. **E-F)** Plasmid templates for C-terminal target tagging with either AID*/3xHA (E) or 3xV5/AID^F^ (F) degrons. Integration cassettes are amplified by PCR using the designated primer annealing sites and primers containing homology regions for the 3’ end of the target gene. In all panels: pr = promoter; tm – terminator; *bla* – beta lactamase gene encoding ampicillin resistance; ori – *E. coli* origin of replication.

The AID system depends on the ability of plant Tir1 to interact with the endogenous core SCF ubiquitin ligase of the target organism, and on the availability of tools to 1) genomically tag the gene of interest with an ABD sequence and 2) stably express the plant *TIR1* gene. To date, inducible target protein degradation using the AID system has been validated in several fungal, protozoan, and metazoan species, including the research model systems *S. cerevisiae* (Nishimura et al., 2009)*, S. pombe* (Watson et al., 2021)*, C. elegans* (Zhang et al., 2015), *D. melanogaster* (Trost et al., 2016), mammalian oocytes (Camlin and Evans, 2019), human cell culture (Holland et al., 2012), and mice (Yesbolatova et al., 2020) and the protozoan pathogens *Toxoplasma gondii* (Brown et al., 2018) and *Plasmodium falciparum* (Kreidenweiss et al., 2013). A limitation of the original AID system is the relatively high concentration of auxin required to induce degradation (typically high µM to low mM), which has been shown to have physiological effects (Nicastro et al., 2021; Prusty et al., 2004) or toxicity (Yesbolatova et al., 2020) in some systems. A recently developed second generation AID system, named AID2, greatly reduces the potential for toxicity or non-specific physiological consequences. AID2 was inspired by the structure-guided engineering of Tir1 to bind the larger, synthetic auxin analogs 5-phenyl-IAA and 5-adamantyl-IAA in plants (Uchida et al., 2018). Replacing wild-type Tir1 with the engineered mutant in AID systems provided efficient target degradation at synthetic auxin concentrations orders of magnitude lower than natural IAA (Nishimura et al., 2020; Yesbolatova et al., 2020).

IPD systems, including AID, have not been established in human fungal pathogens. These systems would provide valuable tools for functional characterization of proteins to better understand pathogen biology, virulence mechanisms, drug resistance, and other clinically relevant processes. Moreover, they could be valuable tools for antifungal drug development. Here, we demonstrate that the AID system works efficiently and rapidly in the human pathogenic yeasts *Candida albicans* and *Candida glabrata.* We have developed reagents for using the original AID and the second generation AID2 systems in lab strains, and provide a blueprint for expanding this system to clinical isolates and other pathogen species.

## RESULTS & DISCUSSION

### Engineering of AID and AID2 systems for use in *C. albicans* and *C. glabrata*

We designed *OsTIR1* integration vectors to be used in common lab strains of *C. albicans* and *C. glabrata* to assess performance of the AID system in these pathogens. For *C. albicans* we synthesized a codon-optimized *OsTIR1* gene with 3xMyc epitope tag and constructed a plasmid with a restriction enzyme-excisable cassette for integration at the *NEUT5L* locus (Gerami-Nejad et al., 2013) with *URA3* selectable marker and *OsTIR1* expression driven by the *ACT1* promoter (**Figure 1B**). We also designed a recyclable, PCR-amplifiable *OsTIR1* integration cassette with *URA3* marker using a previously developed Cre-lox-based *C. albicans* vector system (Dueñas-Santero et al., 2019), in which *OsTIR1* is expressed from the *TDH3* promoter **(Figure 1C).** For *C. glabrata* we constructed a plasmid with excisable cassette for integration at the *TRP1* locus with *LEU2* selectable marker and *OsTIR1-9xMyc* expressed from the *ADH1* promoter (**Figure 1D**). We then introduced the F74A codon change in the *OsTIR1* coding sequence of all plasmids to allow use of the synthetic auxin analog 5-adamantyl IAA and the AID2 system (Nishimura et al., 2020; Yesbolatova et al., 2020).

We designed integration cassettes for C-terminal degron tagging of target proteins for use in both *C. albicans* and *C. glabrata* and inserted them into plasmid backbones to serve as templates for PCR amplification. One cassette contains the coding sequence for the original full 229 amino acid degron domain of the *A. thaliana* IAA17 auxin-binding protein (Nishimura et al., 2009), hereby designated AID^F^, fused to a 3xV5 epitope. The second contains a smaller truncation of the IAA17 degron region (aa 71-114), previously named AID* (Morawska and Ulrich, 2013) fused to a 3xHA epitope. In each case, the degron is followed by a stop codon, transcriptional terminator, and CTG clade codon-optimized *SAT1* marker gene (Reuß et al., 2004) providing nourseothricin resistance for positive selection (**Figure 1E-F**). AID strain construction with these vectors requires two genome integration steps: 1) integration of the *OsTIR1* expression cassette generated by restriction digest or PCR amplification, and 2) integration of a PCR-amplified degron tag at the 3’ end of the desired target gene. For diploid species like *Candida albicans* both alleles can be tagged, or one can be deleted.

### Both AID and AID2 systems provide robust and rapid target degradation in *C. albicans* and *C. glabrata*

For initial testing and comparison of the AID and AID2 systems in *C. albicans*, we tagged the Cdc14 phosphatase at its C-terminus in strains expressing either wild-type OsTir1 or OsTir1^F74A^. Reductions in Cdc14 activity render *C. albicans* cells hypersensitive to cell wall stress and impair hyphal development, allowing facile phenotypic monitoring of AID system performance (Clemente-Blanco et al., 2006; Milholland et al., 2023). We first measured Cdc14-3xV5/AID^F^ degradation by immunoblotting after 60-minute treatment with varying concentrations of IAA in strains expressing wild-type OsTir1, or the synthetic auxin 5-Ad-IAA in strains expressing OsTir1^F74A^. In both cases, Cdc14 level was consistently reduced to ∼1% of its untreated steady-state level (**Figure 2A-B and Figure S1A-B**). With the original AID system, achieving >95% reduction in Cdc14 level required 500 µM IAA, whereas with the AID2 system >95% reduction was achieved at 100,000-fold lower auxin concentration (5 nM 5-Ad-IAA). Using the AID2 system we then compared the effectiveness of the larger 3xV5/AID^F^ degron and the smaller AID*/3xHA degron. Maximal Cdc14 reduction and sensitivity to 5-Ad-IAA concentration were similar with the two degrons (**Figure 2B-C and Figure S1B-C**). Next, we measured the kinetics of Cdc14 degradation in *C. albicans* treated with auxin concentrations that reduced Cdc14 level >95%. The kinetics for the AID and AID2 systems and for the two different degron tags were similar, with half-lives ranging from 5 to 15 min (**Figure 2D-F and Figure S1D-F**).

**Figure 2.**
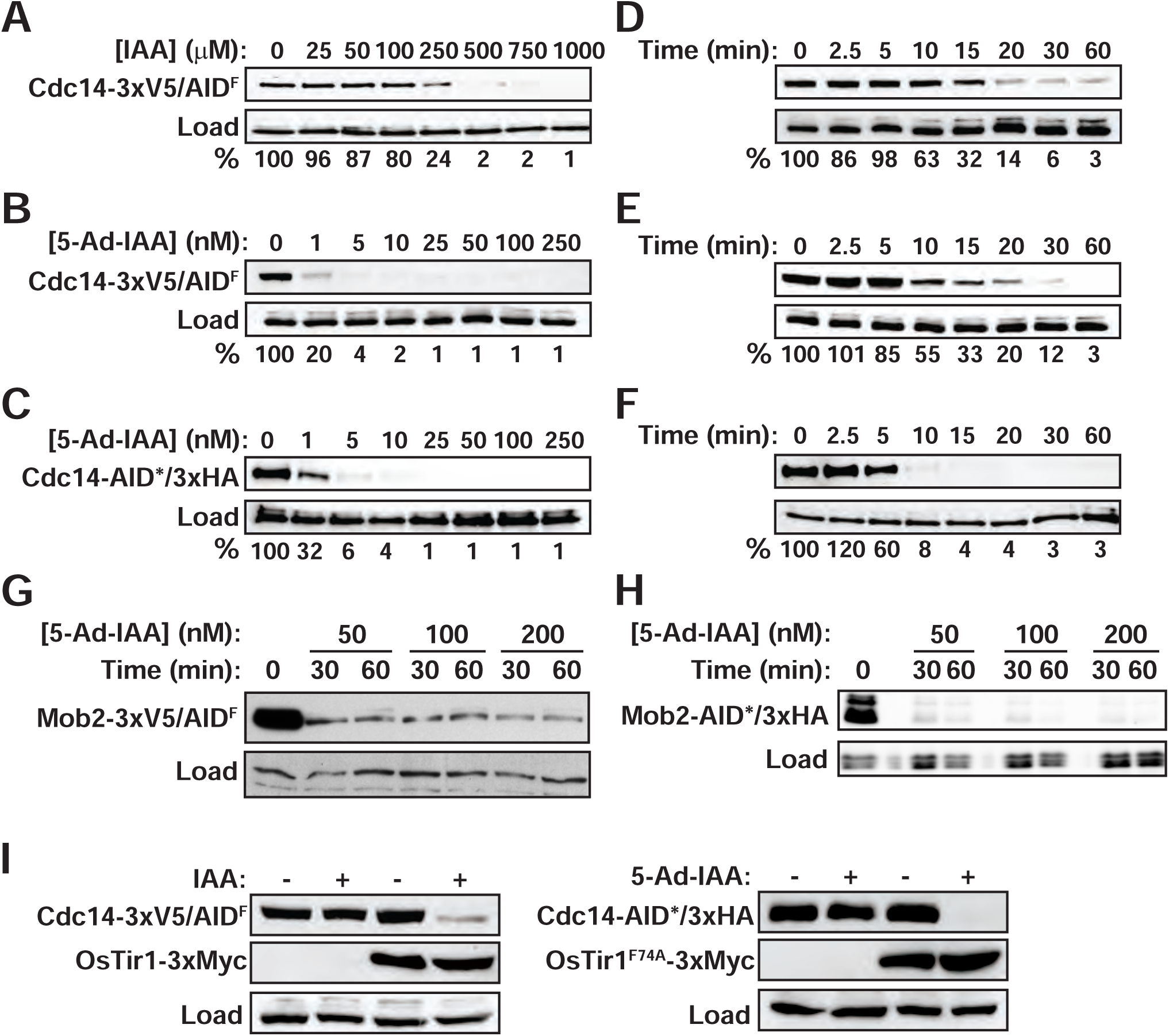
AID technology provides rapid, efficient target degradation in *C. albicans*. **A)** Cdc14-3xV5/AID^F^ degradation in cells expressing wild-type OsTir1 was measured by anti-V5 immunoblotting after treatment of log-phase liquid YPD cultures with the indicated IAA concentrations for 60 min. Anti-PSTAIR antibody was used as a loading control. Percent protein remaining relative to the untreated culture was quantified by digital imaging. **B)** Same as panel A using cells expressing OsTir1^F74A^ and treated with varying 5-Ad-IAA concentrations. **C)** Same as panel B measuring degradation of Cdc14-AID*/3xHA with anti-HA antibody. **D)** Time-dependence of Cdc14-3xV5/AID^F^ degradation in cells expressing wild-type OsTir1 was measured by anti-V5 immunoblotting after treatment of log phase liquid YPD cultures with 500 µM IAA. Percent protein remaining relative to time=0 was quantified by digital imaging. **E)** Same as panel D in cells expressing OsTir1^F74A^ and treated with 50 nM 5-Ad-IAA. **F)** Same as panel E measuring time-dependence of Cdc14-AID*/3xHA degradation using anti-HA antibody. **G)** Degradation of Mob2-3xV5/AID^F^ at the indicated times after 5-Ad-IAA treatment in a strain expressing *OsTIR1^F74A^* from the *TDH3* promoter was monitored by anti-V5 immunoblotting. Anti-Cdc11 was used as a loading control. Anti-PSTAIR was used as a loading control. **H)** Same as A, monitoring degradation of Mob2-AID*/3xHA in a strain expressing *OsTIR1^F74A^* from the *ACT1* promoter. **I)** The dependence of Cdc14-3xV5/AID^F^ degradation on OsTir1 and IAA (left) and of Cdc14-AID*/3xHA degradation on OsTir1^F74A^ and 5-Ad-IAA (right) were determined by immunoblotting with anti-V5 and anti-HA immunoblotting, respectively. Log phase cultures were treated with 500 µM IAA or 50 nM 5-Ad-IAA or mock-treated with an equal volume of DMSO for 60 min prior to harvesting.

We also evaluated the *C. albicans* AID2 system on a different target, the Cbk1 kinase accessory protein, Mob2 (Gutiérrez-Escribano et al., 2011; Song et al., 2008). Mob2-3xV5/AID^F^ was robustly degraded within 30 minutes of 50 nM 5-Ad-IAA addition in a strain expressing *OsTIR1^F74A^* from the *TDH3* promoter, generated with the recyclable *URA3* marker (**Figure 2G**). Mob2-AID*/3xHA was degraded with equal success in a strain expressing *OsTIR1^F74A^* from the *ACT1* promoter (**Figure 2H**). The extent of Mob2 degradation was comparable with the two *OsTIR1^F74A^* expression cassettes.

Finally, we measured the dependence of Cdc14-3xV5/AID^F^ stability on the presence of both OsTir1 and auxin, as auxin-independent target degradation has been reported in some systems (Morawska and Ulrich, 2013; Sathyan et al., 2019; Yesbolatova et al., 2020). We did not observe auxin-independent target degradation in *C. albicans.* Detectable Cdc14 degradation required the presence of both OsTir1 and auxin in both the AID and AID2 systems (**Figure 2I**).

Similar results were observed when AID and AID2 systems were tested in *C. glabrata*. We tagged the histone acetyltransferase, Gcn5, and Cdc14 phosphatase with an AID*/9xMyc degron (Morawska and Ulrich, 2013) in a strain expressing wild-type OsTir1. We also tagged Gcn5 in a strain expressing the AID2 variant OsTir1^F74A^. Steady-state Gcn5-AID*/9xMyc was reduced to ∼1% of the initial levels in log phase cultures, similar to Cdc14 in *C. albicans* (**Figure 3A-B and Figure S2A-B**). >95% reduction required 50 µM IAA in the presence of wild-type OsTir1 and 1 nM 5-Ad-IAA in the presence of OsTir1^F74A^. The kinetics of degradation after treatment an auxin concentration sufficient for >95% degradation were similar in the AID and AID2 systems and slightly faster than for Cdc14 in *C. albicans* (**Figure 3C-D and Figure S2C-D**). Cdc14-AID*/9xMyc degradation in *C. glabrata* was also rapid and efficient, requiring slightly lower IAA concentration than Cdc14-AID*/3xHA in *C. albicans* and exhibiting similar kinetics (**Figure 3E-F).** Similar to *C. albicans*, detectable degradation of Gcn5-AID*/9xMyc in *C. glabrata* was dependent on the presence of both Tir1 and auxin (**Figure 3G**).

**Figure 3.**
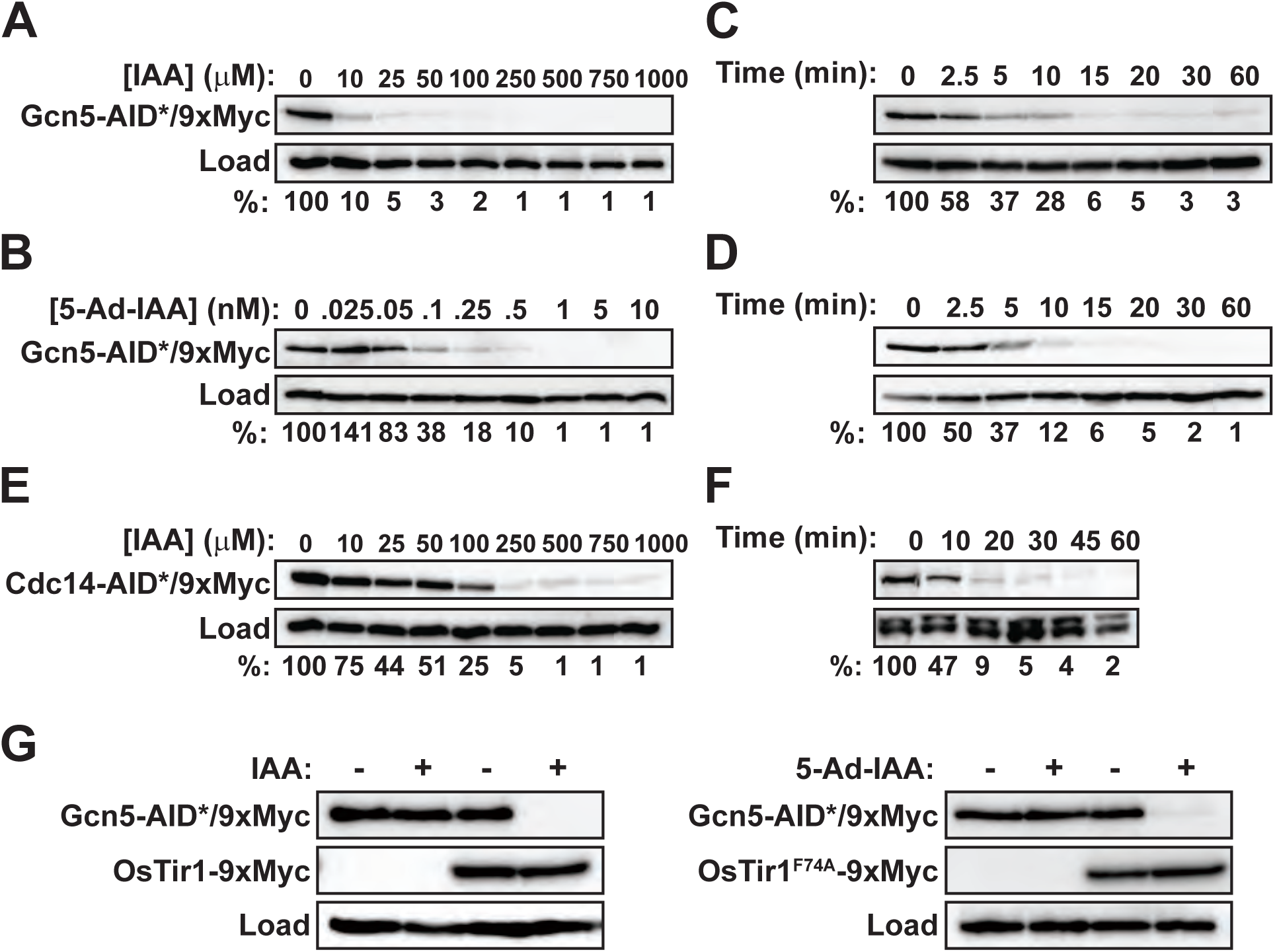
AID technology provides rapid, efficient target degradation in *C. glabrata*. **A)** Gcn5-AID*/9xMyc degradation in cells expressing wild-type OsTir1 was measured by anti-Myc immunoblotting after treatment of log-phase liquid SC cultures with the indicated IAA concentrations for 60 min. Histone H3 antibody was used as a loading control. Percent protein remaining relative to the untreated culture was quantified by digital imaging. **B)** Same as panel A, measuring Gcn5-AID*/9xMyc degradation in cells expressing OsTir1^F74A^ after treatment with the indicated concentrations of 5-Ad-IAA. **C)** Time-dependence of Gcn5-AID*/9xMyc degradation in cells expressing wild-type OsTir1 was measured by anti-Myc immunoblotting after treatment of log phase liquid SC cultures with 100 µM IAA. Percent protein remaining relative to time=0 was quantified by digital imaging. **D)** Same as panel D in cells expressing OsTir1^F74A^ and treated with 5 nM 5-Ad-IAA. **E)** Same as panel A measuring Cdc14-AID*/9xMyc degradation in cells grown in YPD and expressing wild-type OsTir1. **F)** Same as panel C measuring time-dependence of Cdc14-AID*/9xMyc degradation in cells grown in YPD and expressing wild-type OsTir1. **G)** The dependence of Gcn5-AID*/9xMyc degradation on OsTir1 and IAA (left) and on OsTir1^F74A^ and 5-Ad-IAA (right) were determined by anti-Myc immunoblotting. Log phase cultures were treated with 100 µM IAA or 5 nM 5-Ad-IAA or mock-treated with an equal volume of DMSO for 60 min prior to harvesting.

We conclude that the original AID system and the new AID2 system both work effectively to achieve rapid and near-complete target protein loss in *C. albicans* and *C. glabrata* and are therefore likely to be useful tools for protein functional characterization in these species.

### IAA and 5-Ad-IAA have minimal impact on *C. albicans* and *C. glabrata* physiology

Natural auxin can impact cellular physiology in some systems, including *S. cerevisiae* (Nicastro et al., 2021; Prusty et al., 2004) and mice (Yesbolatova et al., 2020). We therefore assessed IAA and 5-Ad-IAA effects on *C. albicans* growth rate, sensitivity to diverse stress conditions, and hyphal differentiation. For the growth assays we used 1 mM IAA and 1 µM 5-Ad-IAA concentrations, well above the concentrations needed for efficient target degradation. The growth of *C. albicans* was unaffected by the presence of IAA or 5-Ad-IAA (**Figure S3A**). In serial dilution spotting assays on YPD agar plates supplemented with oxidative (H_2_O_2_), genotoxic (MMS), or osmotic (NaCl) stresses, or azole antifungal drugs, the presence of IAA and 5-Ad-IAA had no detectable effect on *C. albicans* cell viability or growth rate (**Figure S3B**). Finally, 50 nM 5-Ad-IAA had no impact on *C. albicans* hyphal development induced by serum at 37 °C in liquid cultures (**Figure S3C**). These results indicate that IAA and 5-Ad-IAA are mostly innocuous to *C. albicans*. The very low concentration of 5-Ad-IAA required for target degradation relative to IAA makes it a particularly attractive option for minimizing non-specific physiological effects.

### AID2 phenocopies gene deletions in *C. albicans* and *C. glabrata*

For AID to be useful, target degradation should be extensive enough to induce loss of function phenotypes similar to gene deletions. Loss of Cdc14 function renders *C. albicans* hypersensitive to cell wall stresses, including echinocandin drugs, and also prevents hyphal development on agar plates (Milholland et al., 2023). We used these phenotypes to compare the effects of AID2-mediated Cdc14 depletion to permanent loss-of-function mutations, including homozygous *CDC14* gene deletion and the catalytically-impaired *cdc14^hm^*hypomorphic allele (Milholland et al., 2023). Inclusion of 25 nM 5-Ad-IAA alone in YPD agar plates reduced the growth and viability of *C. albicans CDC14-3xV5/AID^F^* similar to *cdc14Δ/Δ,* demonstrating that AID works efficiently in solid media as well as liquid cultures (**Figure 4A**). Supplementing YPD agar with 50 ng/ml micafungin alone had no impact on *CDC14-3xV5/AID^F^*cells compared to wild-type, but was lethal to *cdc14^hm^/Δ* and *cdc14Δ/Δ*, demonstrating that the AID* degron tag does not significantly compromise Cdc14 function. Importantly, exposing *CDC14-3xV5/AID^F^* cells to just 5 nM 5-Ad-IAA in the presence of 50 ng/ml micafungin severely impaired growth, and 25 nM 5-Ad-IAA with micafungin eliminated growth completely.

**Figure 4.**
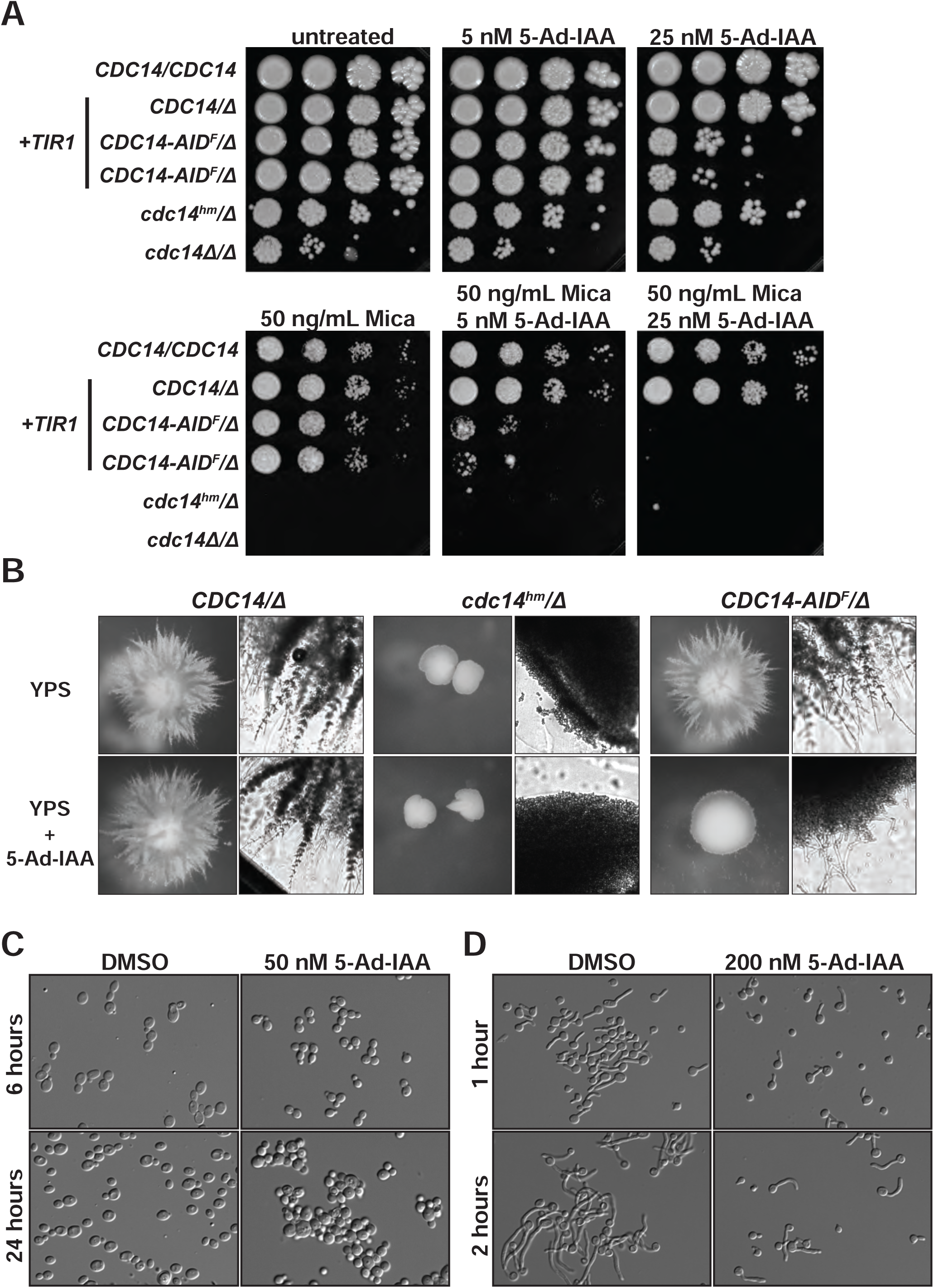
AID effectively phenocopies loss-of-function mutations in *C. albicans*. **A)** Liquid cultures of *C. albicans* strains with the indicated *TIR1* and *CDC14* genotypes were serially diluted and spotted on YPD agar plates supplemented with micafungin and/or auxin, as indicated. Plates were grown at 30 °C for 3 days prior to imaging. The two *CDC14-AID*/Δ* samples are independent transformants from the degron tag integration. The *cdc14^hm^*allele is our previously characterized hypomorphic mutant with reduced catalytic activity (Milholland et al., 2022). **B)** The indicated *C. albicans* strains were grown embedded in YPS agar with or without 150 nM 5-Ad-IAA at 30 °C for 4 days and colonies imaged with a dissecting scope (left) or Cytation 1 imaging plate reader with 4x brightfield objective (right). Fields of view for each imaging method are equivalent across all strains. **C)** DIC images of exponentially growing *MOB2-3xV5/AID^F^*cells expressing *OsTIR1^F74A^* from the *TDH3* promoter (OL3372) incubated in YPD with or without 50 nM 5-Ad-IAA at 28°C for 6h and 24h. **D)** DIC images of *MOB2-AID*/3xHA* cells expressing *OsTIR1^F74A^* from the *ACT1* promoter (OL3309) grown under hypha-inducing conditions (YPD+10% serum at 37°C) in the presence or absence of 50 nM 5-Ad-IAA.

Growth of *C. albicans* embedded within YPS agar medium, or on the surface of Spider agar medium, results in extensive hyphal development, leading to a filamentous colony morphology (**Figure 4B and Figure S4**). Reduced Cdc14 function completely prevents growth of radial hyphae both in embedded YPS agar and on Spider agar. *CDC14-3xV5/AID^F^* cells formed filamentous colonies like wild-type *CDC14* strains on both plate types in the absence of auxin. In contrast, supplementation of YPS or Spider plates with 5-Ad-IAA severely impaired hyphal filament development, similar to *cdc14^hm^/Δ* and *cdc14Δ/Δ*. Deletion of *MOB2* results in a cell separation defect in *C. albicans* (Gutiérrez-Escribano et al., 2011; Song et al., 2008). Including 5-Ad-IAA in liquid *MOB2-3xV5/AID^F^* cultures caused the same cell separation failure reported for *mob2Δ* cells (**Figure 4C).** Cbk1-Mob2 activity is required for maintenance of hyphal growth (Gutiérrez-Escribano et al., 2011; Song et al., 2008). Inclusion of 5-Ad-IAA in YPD-serum liquid medium at 37 °C, conditions that strongly induce hyphae, compromised maintenance of hyphal growth in *MOB2-AID*/3xHA* cells, consistent with loss of Mob2 function (**Figure 4D**).

*C. glabrata gcn5Δ* cells exhibit sensitivity to azole antifungal drugs (Yu et al., 2022), for example in serial dilution plate spotting assays (**Figure 5A**). *GCN5-AID*/9xMyc* cells were indistinguishable from untagged wild-type cells in the presence of fluconazole, confirming that the degron tag does not significantly impair Gcn5 function. In contrast, supplementation of fluconazole with 100 µM IAA in cells expressing wild-type *OsTIR1,* or 5 nM 5-Ad-IAA in cells expressing *OsTIR1^F74A^,* impaired growth of *GCN5-AID*/3xHA* cells, consistent with reduced Gcn5 function. Interestingly, plating *CgCDC14-AID*/9xMyc* cells expressing wild-type OsTir1 on YPD plates containing 500 µM IAA completely prevented growth, suggesting that *CDC14* may be an essential gene in *C. glabrata*, as it is in *S. cerevisiae* (**Figure 5B**). Consistent with this, we have been unable to recover *cdc14Δ* transformants of *C. glabrata* using conventional methods. This highlights a key advantage of the AID system in studying essential genes. Collectively, our results demonstrate that the AID system in *C. albicans* and *C. glabrata* is robust enough to mimic gene deletion phenotypes and should be a useful tool for rapid target protein inactivation to allow phenotypic observation and functional characterization.

**Figure 5:**
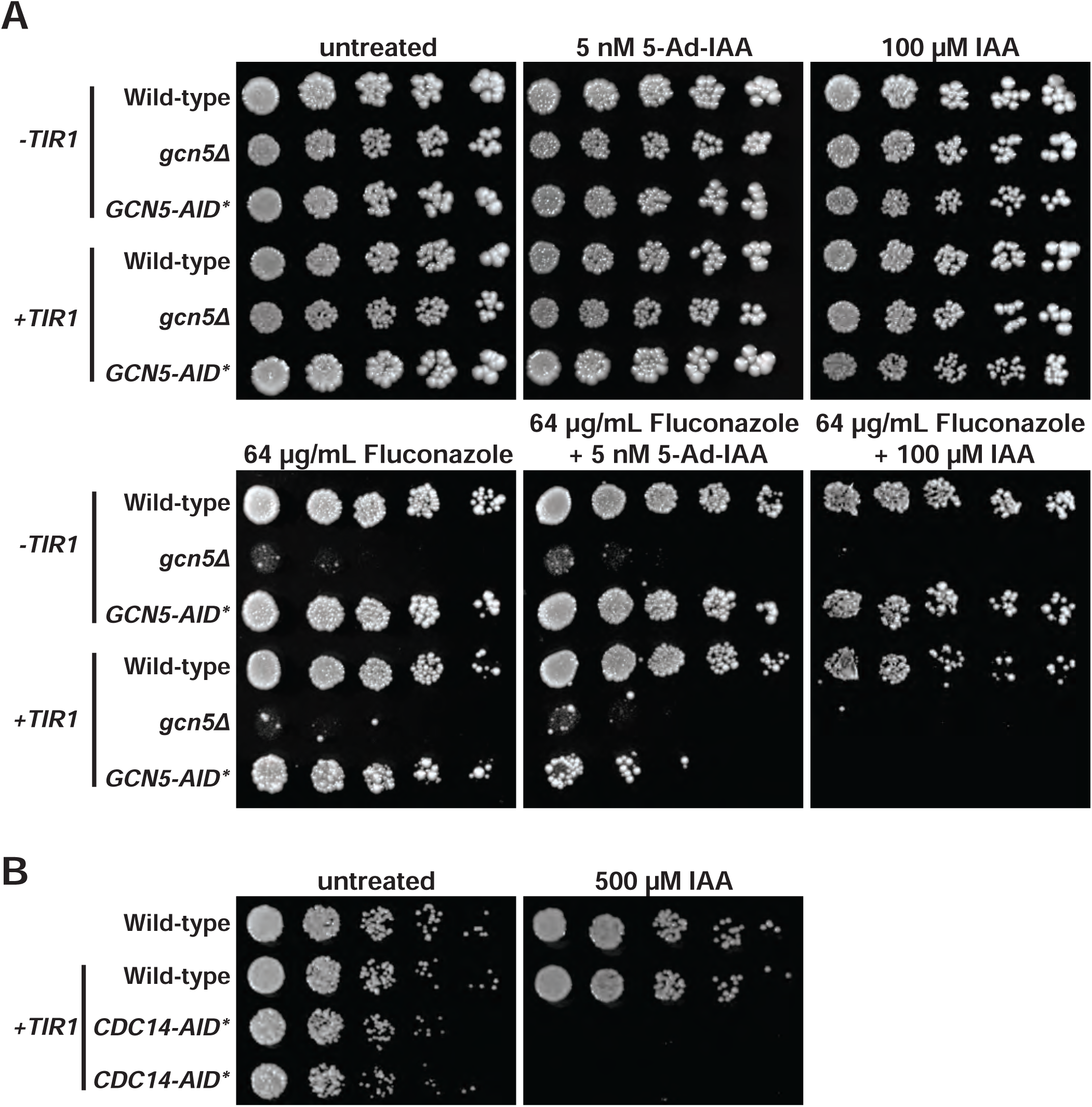
AID effectively phenocopies loss-of-function mutations in *C. glabrata*. **A)** Liquid cultures of the indicated *C. glabrata* strains, with and without integrated *OsTIR1*, were serially diluted and spotted on synthetic complete agar plates supplemented with either 5 nM 5-Ad-IAA (*OsTIR1^F74A^*background) or 100 µM IAA (wild-type *OsTIR1* background) with or without 64 µg/ml fluconazole and grown for 2 days at 30 °C. Note that untreated and fluconazole alone results are shown for the *OsTIR1^F74A^* background strain but were indistinguishable for the wild-type *OsTIR1* strain. The wild-type strain is KKY2001. **B)** Serial dilution spotting assay of the indicated strains on YPD with or without addition of 500 µM IAA. Two independent isolates of the *CgCDC14-AID*/9xMyc* strain were evaluated. Plates were grown for 2 days at 30 °C prior to imaging.

### Conclusions and guidelines

In this study we demonstrated that AID can be successfully implemented in *Candida albicans* and *Candida glabrata* for the rapid and efficient degradation of diverse target proteins. The extent and speed of target protein degradation should make AID a useful functional tool for protein characterization in these species. We expect the system will be readily adaptable to other fungal pathogens as well. AID offers several advantages over other common methods for protein functional characterization. Importantly, AID ensures normal system physiology until auxin is added, avoiding indirect effects of permanent genetic alterations like gene deletions, and longer-term transcriptional repression or RNAi systems. Moreover, AID maintains natural promoter control of the target gene, making it attractive for studying genes with highly regulated expression. The speed at which AID elicits target degradation makes it suitable for studying dynamic processes like signaling pathways, and for mimicking the effects of drugs. With the single requirement of exposing cells to auxin, AID is compatible with diverse experimental conditions, perhaps even live animal models of fungal infections in the future.

The plasmid reagents generated in this study were designed for testing and using AID technology in common lab strains of *C. albicans* and *C. glabrata* and are suitable for experimental use in any strains with the appropriate auxotrophic mutations, or in which the required auxotrophic mutations can be conveniently generated. To make AID useful for studying protein function in clinically-derived pathogen strains and non-model species like the emergent *C. auris*, in which auxotrophic selection is not readily available, we are currently working on expansion of our system to add recyclable antibiotic resistance markers for strain engineering. This may also make the system suitable for use in animal infection models where auxotrophic marker mutations can impact virulence (Kirsch and Whitney, 1991; Lay et al., 1998). The addition of N-terminal degron tag cassettes for targets not compatible with C-terminal fusions, will be an important goal for future development as well.

The three target proteins we selected for testing AID technology behaved ideally, with consistent, extensive, and fast degradation observed specifically after auxin addition only in strains expressing OsTir1. Nonetheless, for each new target protein it is prudent to optimize the auxin concentration, determine the kinetics and maximal percent degradation, and confirm the dependence of degradation on both Tir1 and auxin under the desired experimental conditions. As in other species, AID performance can vary with different target proteins because it functions by changing the equilibrium between protein synthesis and degradation. Differences in protein synthesis rates and mechanisms for regulating steady-state protein levels can result in different AID degradation kinetics and depletion levels. Researchers must also be aware that a degron tag may impair protein function or may not be accessible to OsTir1 and SCF due to protein structure or cellular localization effects. Modifications to AID to overcome problems with inefficient, or auxin-independent degradation have been designed and validated in model organisms (Li et al., 2019; Mendoza-Ochoa et al., 2019; Sathyan et al., 2019; Yesbolatova et al., 2019) and may be feasible to implement in *Candida* species if needed.

## METHODS

### Plasmid construction

Plasmids constructed in this study are listed in Table S1. All pHLP plasmids were created using the In-Fusion cloning system (Takara Biosciences) and were confirmed by Wide-seq analysis. CTG clade codon-optimized coding sequences for 1) *OsTIR1* fused to 3xMyc, and 2) *3xV5/AID^F^* were synthesized by Twist Bioscience. pDIS3 (Gerami-Nejad et al., 2013), which has the targeting sequences for integration at the NEUT5L locus, was used as the starting point for pHLP710 assembly. An XhoI fragment containing the SAT1 marker was excised from pDIS3 and replaced with PCR-amplified *C. albicans URA3* from p347 (Milholland et al., 2023), *C. albicans ACT1* promoter and intron cassette from pSFS2 (Reuß et al., 2004), and codon-optimized synthetic *OsTIR1-3xMyc.* The In-Fusion assembly was designed so the existing *TEF* terminator from pDIS3 followed the *OsTIR1-3xMyc* coding sequence. Site-directed mutagenesis (QuikChange II, Agilent) was used to introduce the Phe74 to Ala74 codon change in the *OsTIR1* coding sequence of pHLP710, creating pHLP712 for the AID2 system (Nishimura et al., 2020; Yesbolatova et al., 2020). To create template plasmids for amplification of C-terminal degron tagging cassettes with the *SAT1* selectable marker we excised the existing degron/marker region from the original AID template plasmid pAR1070 (Powers and Hall, 2017) by digestion with NotI and HindIII and used In-Fusion assembly to insert 1) either the synthetic 3xV5/AID^F^ coding sequence (pHLP701) or AID*/3xHA sequence (pHLP700) amplified from pHyg-AID*-6HA (Morawska and Ulrich, 2013) followed by 2) the *S. cerevisiae ADH1* terminator and 3) the *ACT1* promoter-*SAT1* cassette with *C. albicans URA3* terminator amplified from pSFS2.

Plasmids pJC608 and pJC609 are based on the pFA-Clox plasmid toolkit (Dueñas-Santero et al., 2019) and were constructed using the NEBuilder HiFi DNA Assembly Cloning Kit (New England Biolabs) following manufacturer’s instructions. Primer design for PCR amplification of the different modules was performed with the NEBuilder Assembly tool (http://nebuilder.neb.com/). First, a 953 bp region from the *TDH3* promoter (−950 to +3) was amplified from genomic DNA and assembled into pFA-URA3-Clox vector digested with ClaI to produce the plasmid pFA-URA3-Clox-TDH3p. Second, a 2120 bp fragment containing the OsTIR1 gene and the TEF1 terminator sequence was amplified from pHLP710 or pHLP712 and assembled into the pFA-URA3-Clox-TDH3p vector linearized with EcoRV to give rise to the pFA-URA3-Clox-TDH3p-OsTIR1 (pJC608) and pFA-URA3-Clox-TDH3p-OsTIR1^F74A^ (pJC609).

The plasmid pBV90 was constructed by first digesting the plasmid pBYP6744 (Tanaka et al., 2015) with EcoRI and XmaI to excise the *ADH1pr-OsTIR1* construct. By Gibson cloning (New England Biolabs), *ADH1pr-OsTIR1* was subsequently added into the pUC19 backbone along with Sc*LEU2* cassette and flanking regions of Cg*TRP1* for targeted insertion. pBV514 was constructed by the same strategy as pBV90 with the exception of the F74A mutation in the *OsTIR1* gene. The F74A mutation was constructed by Gibson cloning with overlapping primers harboring the desired mutation.

### Strain construction

Oligonucleotide primers used for strain constructions are listed in Table S2. All strains created or used in this study are listed in Table S3. For *OsTIR1* integration at *NEUT5L* in *C. albicans,* 25 µg pHLP710 or pHLP712 were digested with *SfiI,* ethanol-precipitated, and transformed by electroporation with selection on synthetic medium lacking uracil. Integration was confirmed by locus-specific PCR and anti-Myc immunoblotting. To integrate the *TDH3p-OsTIR1* cassettes from pJC608 and pJC609 at *NEUT5L* the *URA3*-Clox-*TDH3p-OsTIR1* modules were amplified by PCR with primers S1-NEUT5L and S2-NEUT5L and transformed by electroporation with selection for uracil prototrophy. Elimination of the *URA3* marker was performed as described (Dueñas-Santero et al., 2019). pBV90 and pBV514 were digested with PacI and SbfI and transformed into *C. glabrata* (KKY2001) with selection for leucine prototrophy to make the strains BVGC16 and BVGC612, respectively.

Degron/epitope tags amplified from pHLP700 and pHLP701 were integrated at the 3’ end of the *CDC14* gene using PCR primers containing 21 template annealing bases and 69 bases of homology immediately upstream of the *CDC14* stop codon and downstream of a CRISPR guide RNA recognition site. Guide RNA was selected in the 3’ UTR as close to the stop codon as possible. PCR products (∼1-2 µg) were ethanol-precipitated and resuspended in a small volume of sterile water and transformed by electroporation using a modified CRISPR-Cas9 RNP procedure described previously (Grahl et al., 2017). For tagging Mob2 at the C-terminus, cassettes were amplified from pHLP700 and pHLP701 using PCR primers listed in Table S2. PCR products were ethanol-precipitated and resuspended in 10 µl of sterile water and transformed by electroporation.

### Media and cell culture

Liquid cultures of *C. albicans* and *C. glabrata* were grown in YPD medium (10 g/L yeast extract, 20 g/L peptone, 20 g/L glucose) at 30°C with shaking at 225 rpm. Agar was added to 20% (w/v) for growth on solid medium. For *C. albicans ura3* auxotrophic strains YPD was supplemented with 8 µg/mL uridine. For agar plate spotting assays, single colonies were selected and grown to saturation. Saturated cultures were diluted to a starting OD_600_ = 0.125 and then serially diluted in 8-fold steps, with 5 µL of four consecutive dilutions spotted on plates and grown at 30°C for 3-5 days. *C. albicans cdc14Δ/Δ* cultures were diluted one less time than other strains and spots therefore correspond to 8-fold higher absorbance; this was necessary to normalize colony density on untreated control plates. For microplate growth assays, single colonies were selected and grown to saturation. Saturated cultures were diluted to OD_600_ = 0.02 in YPD and mixed with an equal volume of YPD, YPD + 2 mM IAA, or YPD + 2 µM 5-Ad-IAA in a sterile, 96-well suspension culture plate. Plates were incubated at 30°C with continuous orbital shaking at 425 cpm. OD_600_ was measured every 15 minutes for 24 hours. Hyphal induction under embedded YPS (10 g/L yeast extract, 20 g/L peptone, 20 g/L sucose) agar conditions was assessed as described previously (Mulhern et al., 2006). Hyphal induction was also monitored on Spider media agar plates (10 g/L beef broth, 10 g/L mannitol, 2 g/L K_2_HPO_4_, 20 g/L agar) grown at 37°C for up to 10 days. Auxin and stress agents at the indicated concentrations were added to cooled media immediately prior to pouring plates. For *C. glabrata* spotting assays, strains were inoculated in synthetic complete (SC) medium (Sunrise Science Products) and grown to saturation overnight. Cultures were diluted to an OD_600_ of 0.1 and grown in SC to log phase under shaking at 30°C. Each strain was spotted in five-fold dilutions starting at an OD_600_ of 0.01 on SC plates with or without the indicated concentrations of fluconazole (Cayman Chemical), 5-Ad-IAA, and IAA. Plates were grown at 30°C for 2-3 days and imaged.

For hyphal induction analysis in liquid medium, *C. albicans* strains were grown overnight to saturation in YPD and then diluted to OD_600_=0.5 in YPD + 10% serum supplemented with either 200 nM 5-Ad-IAA or an equal volume of DMSO at 37°C. Samples were collected 1 and 2 hours later and imaged by DIC microscopy.

### SDS-PAGE and immunoblotting

Total protein extracts were prepared as described (Milholland et al., 2022). Proteins were separated on 10% tris-glycine SDS-PAGE gels, transferred to 0.45 µm nitrocellulose membranes (Bio-Rad) and probed overnight at 4°C with mouse anti-HA (1:5,000; Sigma-Aldrich, 12CA5), rabbit anti-V5 (1:5,000; Invitrogen, MA5-15253), mouse anti-c-Myc (1:2,000-1:5,000; Sigma-Aldrich, 9E10), or rabbit anti-PSTAIR (1:5,000; Millipore-Sigma, 06-923). The rabbit polyclonal anti-H3 antibody was generated by Pocono Rabbit Farm & Laboratory and used at 1:100,000 dilution. Secondary anti-mouse and anti-rabbit antibodies conjugated to horseradish peroxidase were from Jackson ImmunoResearch (115-035-003 or 111-035-003) and used at 1:10,000 dilution for 60 min. at 4°C. Immunoblots were developed with Clarity Western ECL Substrate (Bio-Rad, 170-5060) and imaged on a ChemiDoc MP multimode imager (Bio-Rad).

## ACKNOWLEDGEMENTS

SDB is supported by grant AI136995, SM-R by grant AI152494, and MCH, SDB and JC-B by grant AI168050 from the National Institutes of Health, National Institute of Allergy and Infectious Diseases. JBG was also supported by the National Institute of Allergy and Infectious Diseases under award number T32AI148103. The content is solely the responsibility of the authors and does not necessarily represent the official view of the National Institutes of Health. The work was also supported by the Indiana Clinical and Translational Sciences Institute funded, in part by Award Number UL1TR002529 from the National Institutes of Health, National Center for Advancing Translational Sciences, Clinical and Translational Sciences Award. MCH and SDB were supported by an award from the Purdue University AgSEED program. CRVA and JCB are funded by project PID2020-118109RB-I00 from the Spanish MCIN/AEI/10.13039/501100011033 and by Internationalization Project “CL-EI-2021-08-IBFG Unit of Excellence” of the Spanish National Research Council (CSIC), funded by the Regional Government of Castile and Leon and co-financed by the European Regional Development Fund (ERDF “Europe drives our growth”).

## Supplemental Figure Legends

**Figure S1.**
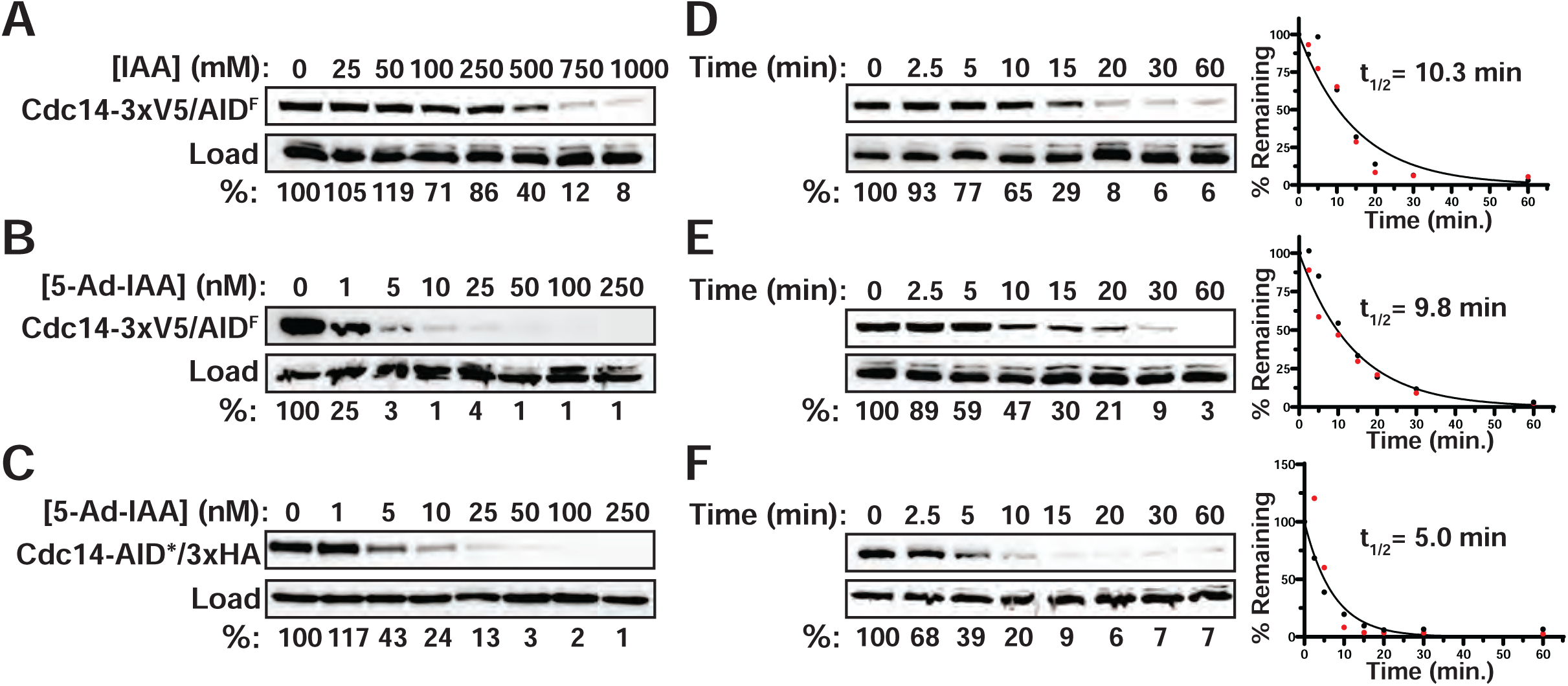
AID technology provides rapid, efficient target degradation in *C. albicans*. **A-C)** Repeat trials for experiments shown in Figure 2 panels A-C. Percent protein remaining relative to the untreated culture was quantified by digital imaging. **D-F)** Repeat trials for Figure 2 panels D-F showing degradation kinetics of Cdc14 proteins in response to IAA (D) or 5-Ad-IAA (E-F) treatment. Percent protein remaining relative to time=0 was quantified by digital imaging. These values, combined with those from Figure 2, were used to calculate half-life from a simple exponential decay function using GraphPad Prism.

**Figure S2.**
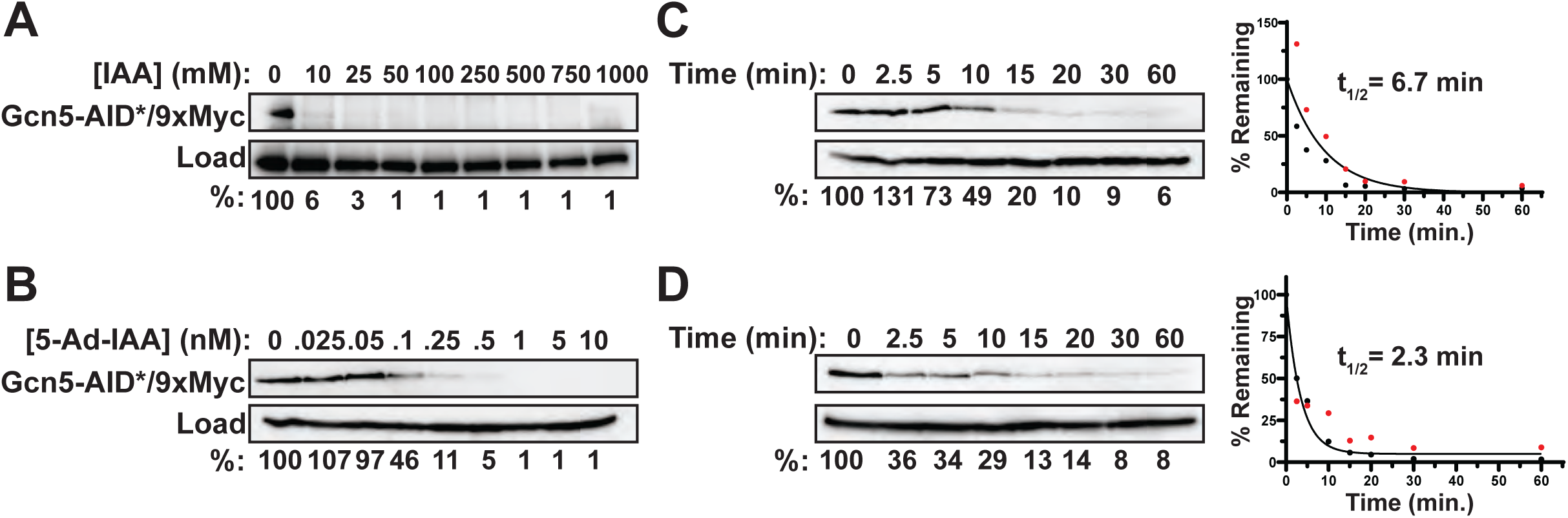
AID technology provides rapid, efficient target degradation in *C. glabrata*. **A-B)** Repeat trials for experiments shown in Figure 4 panels A-B. Percent protein remaining relative to the untreated culture was quantified by digital imaging. **C-D)** Repeat trials for Figure 4 panels C-D showing degradation kinetics of Gcn5 proteins in response to IAA (C) or 5-Ad-IAA (D) treatment. Percent protein remaining relative to time=0 was quantified by digital imaging. These values, combined with those from Figure 4, were used to calculate half-life from a simple exponential decay function using GraphPad Prism.

**Figure S3.**
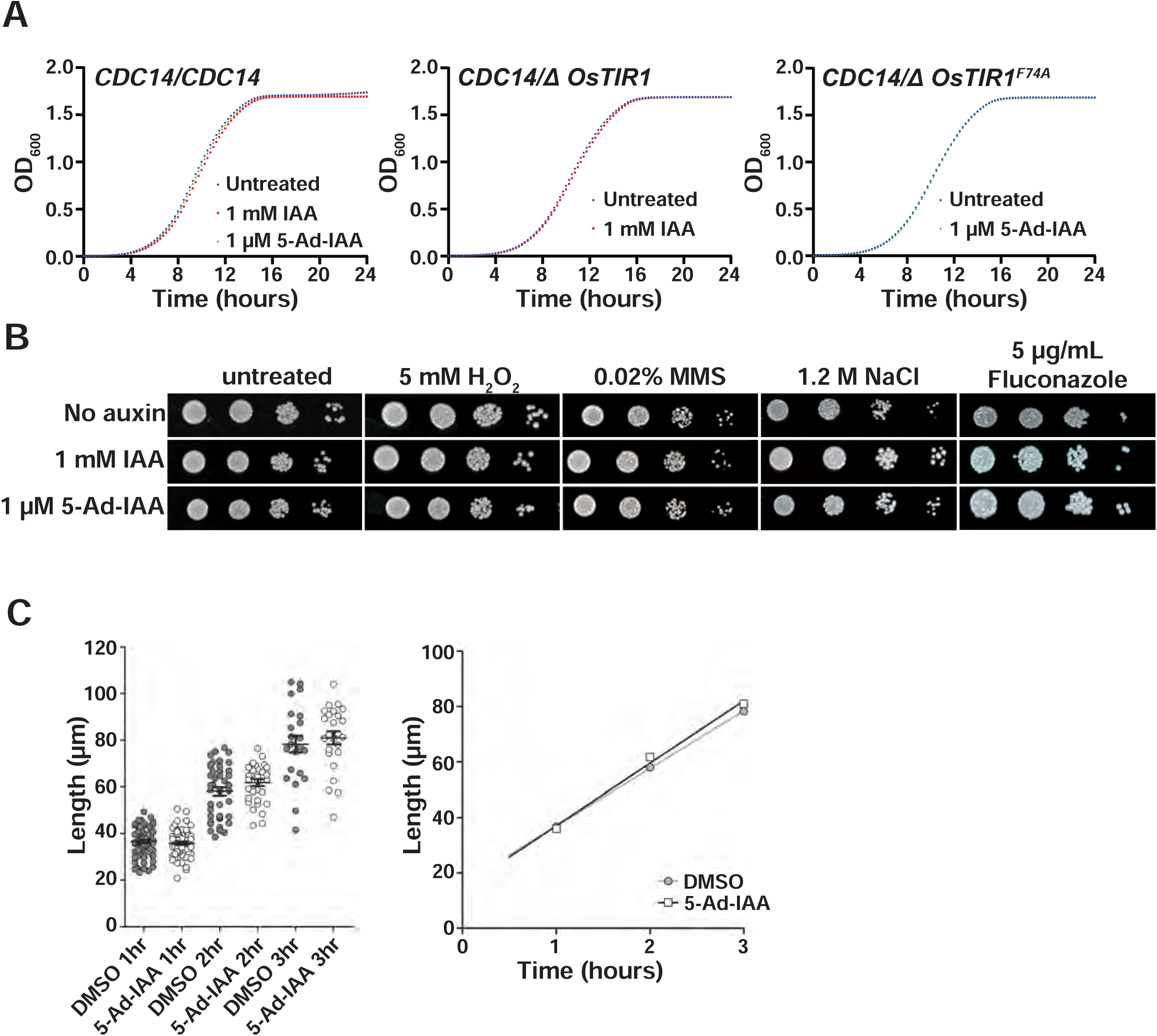
IAA and 5-Ad-IAA do not affect proliferation, stress sensitivity, or hyphal growth of *C. albicans*. **A)** Liquid cultures of *C. albicans* CAI4 (left) or CAI4 with integrated *OsTIR1* (middle) or *OsTIR1^F74A^* (right) were back-diluted to OD_600_ = 0.01 in YPD in a 96-well microplate, supplemented with the indicated concentrations of IAA or 5-Ad-IAA, and grown at 30°C with shaking for 24 hours in a plate reader, measuring absorbance at 600 nm. **B)** A liquid culture of *C. albicans* strain CAI4 was serially diluted and spotted on YPD agar plates supplemented with the indicated chemicals in the absence or presence of 1 mM IAA or 1 µM 5-Ad-IAA and grown at 30 °C for 3 days prior to imaging. Concentrations of H_2_O_2_, MMS, NaCl, and fluconazole were chosen slightly lower than those causing noticeable toxicity compared to the “No Stress” plates. **C)** Liquid cultures of the wild-type BWP17 strain were transferred to YPD + 10% serum at 37 °C to induce hyphal growth in the absence or presence of 50 nM 5-Ad-IAA and hyphal length was measured at several timepoints. Mean lengths were plotted as a function of serum induction time on the right.

**Figure S4.**
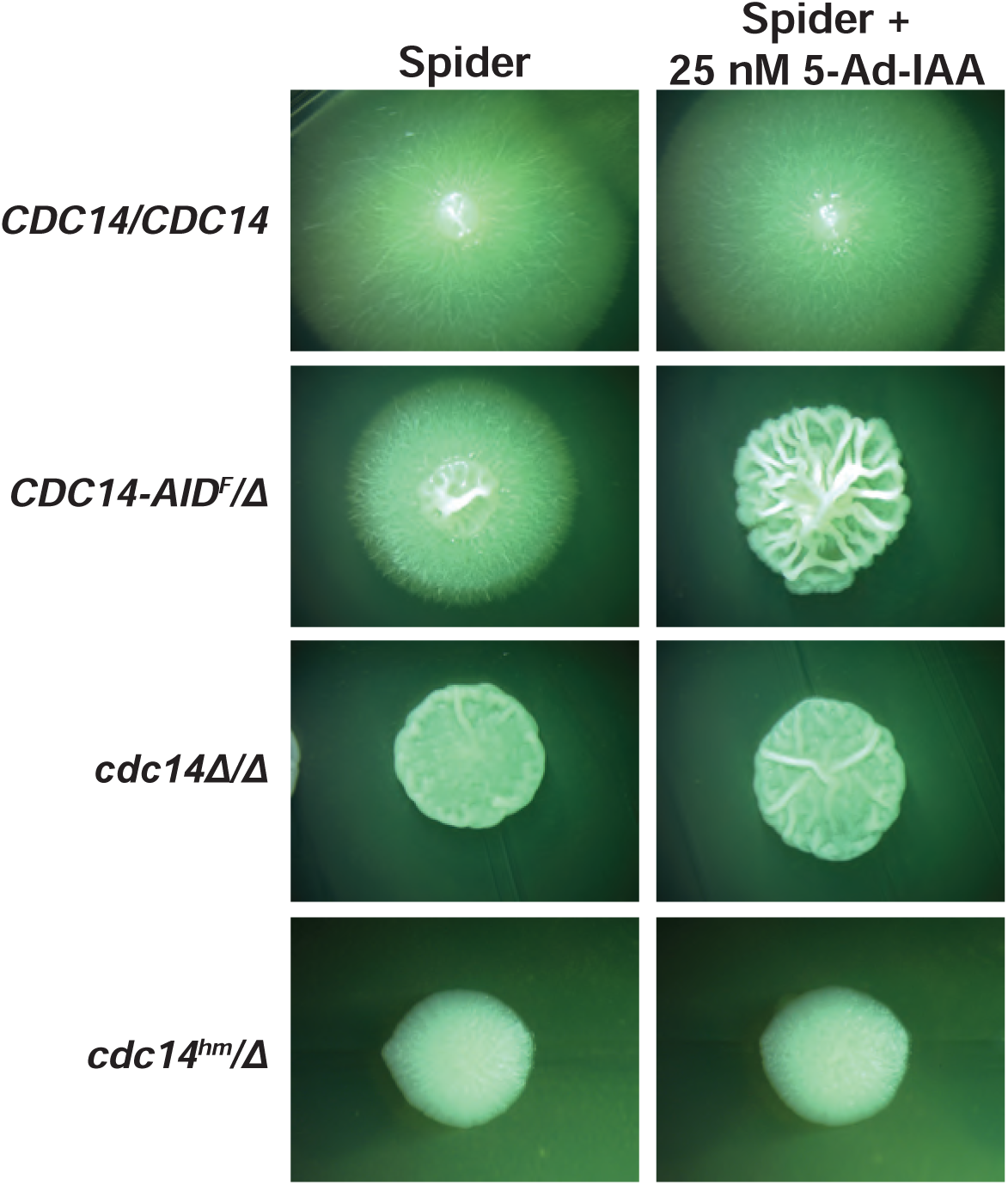
AID effectively phenocopies loss-of-function mutations in *C. albicans*. The indicated *C. albicans* strains were grown as individual colonies on Spider agar plates at 37 °C for 7 days prior to imaging. All images represent identical plate areas. The *CDC14-AID^F^/Δ* strain also has integrated *OsTIR1^F74A^*.

## SUPPLEMENTAL TABLES

**Table S1.** Plasmids created in this study.

| Name | Backbone | Yeast Marker | Expressed gene or tag |
| --- | --- | --- | --- |
| pHLP710 | pDIS3 (Gerami-Nejad et al., 2013) | *URA3* | *ACT1pr-OsTIR1-3xMyc* |
| pHLP712 | pDIS3 (Gerami-Nejad et al., 2013) | *URA3* | *ACT1pr-OsTIR1<sup>F74A</sup>-3xMyc* |
| pHLP700 | pAR1070 (Powers and Hall, 2017) | *SAT1* | AID*-3xHA |
| pHLP701 | pAR1070 (Powers and Hall, 2017) | *SAT1* | 3xV5-AID<sup>F</sup> |
| pBV90 | pUC19 | *LEU2* | *ADH1pr-OsTIR1-9xMyc* |
| pBV514 | pUC19 | *LEU2* | *ADH1pr-OsTIR1<sup>F74A</sup>-9xMyc* |
| pJC608 | pFA-URA3-Clox (Dueñas-Santero et al., 2019) | *URA3* | *TDH3pr-OsTIR1* |
| pJC609 | pFA-URA3-Clox (Dueñas-Santero et al., 2019) | *URA3* | *TDH3pr-OsTIR1<sup>F74A</sup>* |

**Table S2.**
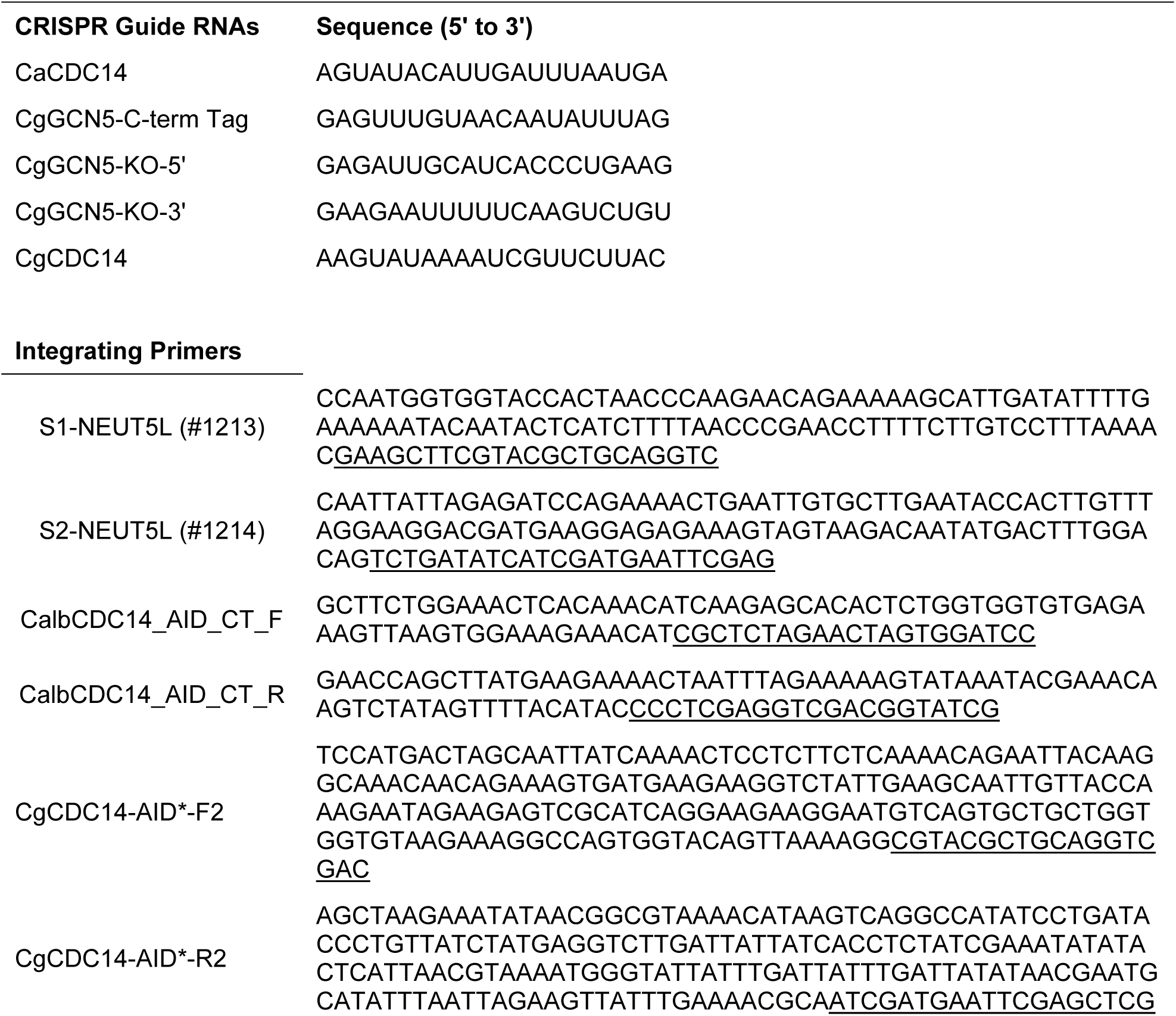
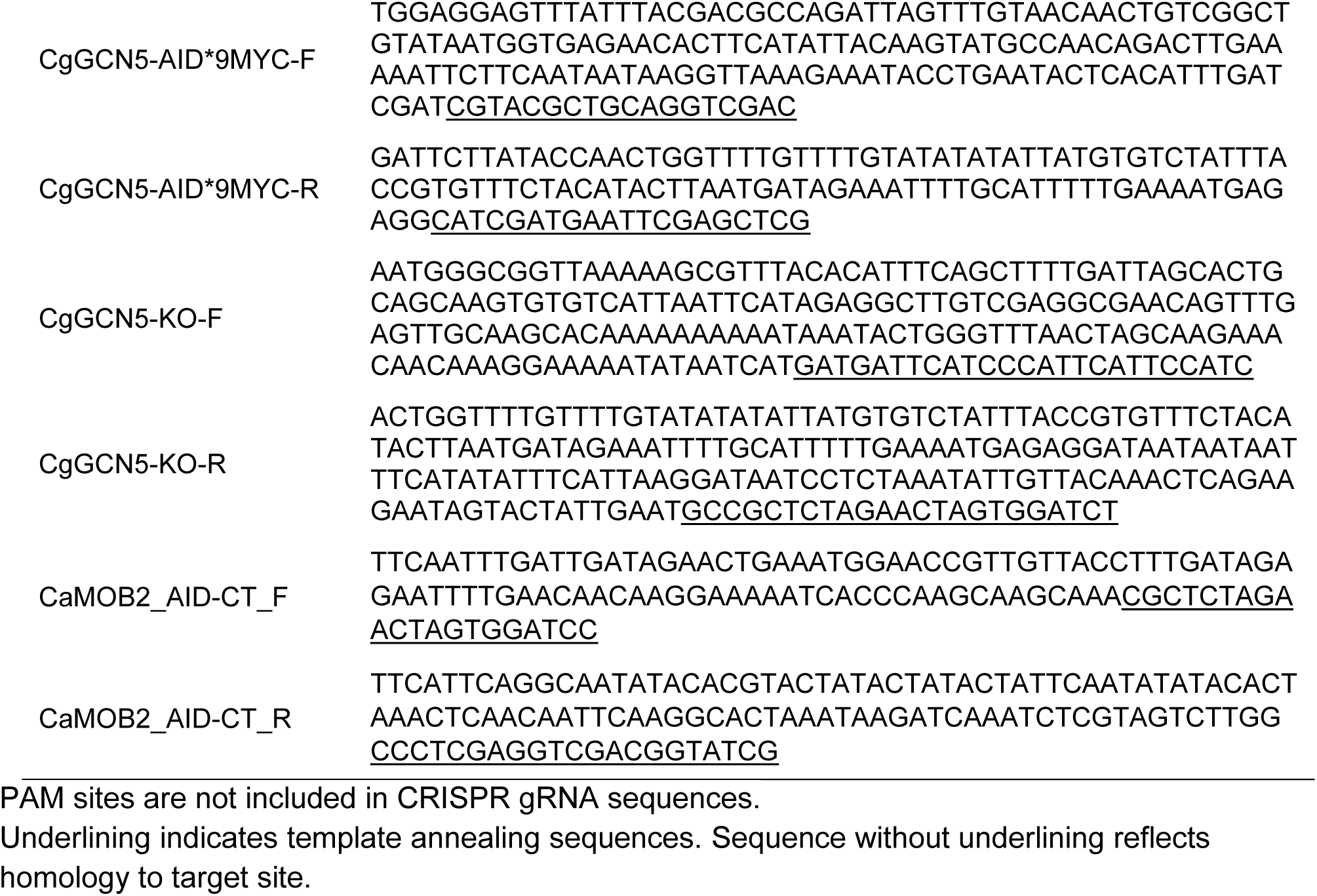
Oligonucleotides for strain engineering.

**Table S3.** Yeast strains.

| Name | Genotype | Reference |
| --- | --- | --- |
| ***C. albicans*** |  |  |
| CAI4 | *ura3::imm434/ura3::imm434* | (Fonzi and Irwin, 1993) |
| BWP17 | *ura3::imm434/ura3::imm434 his1::hisG/his1::his arg4::hisG/arg4::hisG* | (Wilson et al., 1999) |
| JC2711 | *CAI4 NEUT5L::URA3 cdc14::hisG/cdc14::hisG* | (Milholland et al., 2023) |
| JC2712 | *CAI4 NEUT5L::URA3* | (Milholland et al., 2023) |
| JC2721 | *CAI4 CDC14-3xHA:URA3/cdc14::hisG* | (Milholland et al., 2023) |
| HCAL102 | *CAI4 cdc14<sup>(Q414A/P415A/K417A)</sup>-3xHA:URA3/cdc14::hisG* | (Milholland et al., 2023) |
| JC5 | *CAI4 CDC14/cdc14::hisG* | (Clemente-Blanco et al., 2006) |
| HCAL110 | *CAI4 CDC14/cdc14::hisG NEUT5L:OsTIR1-3xMyc:URA3* | This study |
| HCAL111 | *CAI4 CDC14/cdc14::hisG NEUT5L:OsTIR1<sup>F74A</sup>-3xMyc:URA3* | This study |
| HCAL112 | *CAI4 CDC14-3xV5-AID<sup>F</sup>:SAT1/cdc14::hisG* | This study |
| HCAL113 | *CAI4 CDC14-AID\*-3xHA:SAT1/cdc14::hisG* | This study |
| HCAL125 | *CAI4 CDC14-3xV5-AID<sup>F</sup>:SAT1/cdc14::hisG NEUT5L:OsTIR1-3xMyc:URA3* | This study |
| HCAL126 | *CAI4 CDC14-3xV5-AID<sup>F</sup>:SAT1/cdc14::hisG NEUT5L:OsTIR1<sup>F74A</sup>-3xMyc:URA3* | This study |
| HCAL128 | *CAI4 CDC14-AID\*-3xHA:SAT1/cdc14::hisG NEUT5L:OsTIR1<sup>F74A</sup>-3xMyc:URA3* | This study |
| JC2958 | *BWP17 NEUT5L::loxP-TDH3p-OsTIR1-3xMyc* | This study |
| JC2960 | *BWP17 NEUT5L::loxP-TDH3p-OsTIR1<sup>F74A</sup>-3xMyc* | This study |
| OL3372 | *BWP17 mob2::LUL/MOB2-3xV5-AID<sup>F</sup>:SAT1; NEUT5L::loxP-TDH3p-OsTIR1<sup>F74A</sup>-3xMyc* | This study |
| OL3309 | *BWP17 mob2::loxP-MOB2-AID\*-3xHA::SAT1; NEUT5L::URA3-ACT1p-OsTIR1<sup>F74A</sup>-3xMyc* | This study |
| ***C. glabrata*** |  |  |
| KKY2001 | *his3::FRT leu2::FRT trp1::FRT* | (Paul et al., 2018) |
| BVGC16 | *his3::FRT leu2::FRT trp1::ADH1p-OsTIR1-9xMyc:LEU2* | This study |
| BVGC612 | *his3::FRT leu2::FRT trp1::ADH1p-OsTIR1<sup>F74A</sup>-9xMyc:LEU2* | This study |
| SDBY1700 | *his3::FRT leu2::FRT trp1::ADH1p-OsTIR1-9xMyc:LEU2 GCN5-AID\*-9xMyc* | This study |
| SDBY1701 | *his3::FRT leu2::FRT trp1::ADH1p-OsTIR1<sup>F74A</sup>-9xMyc:LEU2 GCN5-AID\*-9xMyc* | This study |
| SDBY1702 | *his3::FRT leu2::FRT trp1::FRT GCN5-AID\*-9xMyc* | This study |
| SDBY1703 | *his3::FRT leu2::FRT trp1::ADH1p-OsTIR1-9xMyc:LEU2 CDC14-AID\*-9xMyc* | This study |
| SDBY1704 | *his3::FRT leu2::FRT trp1::FRT gcn5::SAT1* | This study |
| SDBY1705 | *his3::FRT leu2::FRT trp1::ADH1p-OsTIR1-9xMyc:LEU2 gcn5::SAT1* | This study |
| SDBY1706 | *his3::FRT leu2::FRT trp1::ADH1p-OsTIR1<sup>F74A</sup>-9xMyc:LEU2 gcn5::SAT1* | This study |

## REFERENCES

Bellí, G., Garí, E., Piedrafita, L., Aldea, M., Herrero, E., 1998. An activator/repressor dual system allows tight tetracycline-regulated gene expression in budding yeast. Nucleic Acids Res. 26, 942–947. https://doi.org/10.1093/nar/26.4.942

Brown, K.M., Long, S., Sibley, L.D., 2018. Conditional Knockdown of Proteins Using Auxin-inducible Degron (AID) Fusions in Toxoplasma gondii. Bio-Protoc. 8, e2728. https://doi.org/10.21769/BioProtoc.2728

Camlin, N.J., Evans, J.P., 2019. Auxin-inducible protein degradation as a novel approach for protein depletion and reverse genetic discoveries in mammalian oocytes†. Biol. Reprod. 101, 704–718. https://doi.org/10.1093/biolre/ioz113

Clemens, J.C., Worby, C.A., Simonson-Leff, N., Muda, M., Maehama, T., Hemmings, B.A., Dixon, J.E., 2000. Use of double-stranded RNA interference in Drosophila cell lines to dissect signal transduction pathways. Proc. Natl. Acad. Sci. U. S. A. 97, 6499–6503. https://doi.org/10.1073/pnas.110149597

Clemente-Blanco, A., Gonzalez-Novo, A., Machin, F., Caballero-Lima, D., Aragon, L., Sanchez, M., de Aldana, C.R., Jimenez, J., Correa-Bordes, J., 2006. The Cdc14p phosphatase affects late cell-cycle events and morphogenesis in Candida albicans. J Cell Sci 119, 1130–43. https://doi.org/jcs.02820 [pii] 10.1242/jcs.02820

Dong, G., Ding, Y., He, S., Sheng, C., 2021. Molecular Glues for Targeted Protein Degradation: From Serendipity to Rational Discovery. J. Med. Chem. 64, 10606–10620. https://doi.org/10.1021/acs.jmedchem.1c00895

Dueñas-Santero, E., Santos-Almeida, A., Rojo-Dominguez, P., Rey, F. del, Correa-Bordes, J., Aldana, C.R.V. de, 2019. A new toolkit for gene tagging in Candida albicans containing recyclable markers. PLOS ONE 14, e0219715. https://doi.org/10.1371/journal.pone.0219715

Fonzi, W.A., Irwin, M.Y., 1993. Isogenic strain construction and gene mapping in Candida albicans. Genetics 134, 717–728. https://doi.org/10.1093/genetics/134.3.717

Gerami-Nejad, M., Zacchi, L.F., McClellan, M., Matter, K., Berman, J., 2013. Shuttle vectors for facile gap repair cloning and integration into a neutral locus in Candida albicans. Microbiology 159, 565–579. https://doi.org/10.1099/mic.0.064097-0

Grahl, N., Demers, E.G., Crocker, A.W., Hogan, D.A., 2017. Use of RNA-Protein Complexes for Genome Editing in Non-albicans Candida Species. mSphere 2. https://doi.org/10.1128/mSphere.00218-17

Gutiérrez-Escribano, P., González-Novo, A., Suárez, M.B., Li, C.-R., Wang, Y., de Aldana, C.R.V., Correa-Bordes, J., 2011. CDK-dependent phosphorylation of Mob2 is essential for hyphal development in Candida albicans. Mol. Biol. Cell 22, 2458–2469. https://doi.org/10.1091/mbc.e11-03-0205

Holland, A.J., Fachinetti, D., Han, J.S., Cleveland, D.W., 2012. Inducible, reversible system for the rapid and complete degradation of proteins in mammalian cells. Proc. Natl. Acad. Sci. 109, E3350. https://doi.org/10.1073/pnas.1216880109

Kamath, R.S., Fraser, A.G., Dong, Y., Poulin, G., Durbin, R., Gotta, M., Kanapin, A., Le Bot, N., Moreno, S., Sohrmann, M., Welchman, D.P., Zipperlen, P., Ahringer, J., 2003. Systematic functional analysis of the Caenorhabditis elegans genome using RNAi. Nature 421, 231–237. https://doi.org/10.1038/nature01278

Kirsch, D.R., Whitney, R.R., 1991. Pathogenicity of Candida albicans auxotrophic mutants in experimental infections. Infect. Immun.

Kreidenweiss, A., Hopkins, A.V., Mordmüller, B., 2013. 2A and the Auxin-Based Degron System Facilitate Control of Protein Levels in Plasmodium falciparum. PLOS ONE 8, e78661. https://doi.org/10.1371/journal.pone.0078661

Lay, J., Henry, K.L., Clifford, J., Bulawa, C.E., Becker, J.M., Yigal, K., 1998. Altered Expression of Selectable Marker URA3 in Gene-Disrupted Candida albicans Strains Complicates Interpretation of Virulence Studies. Infect. Immun. 66, 5301–5306. https://doi.org/10.1128/IAI.66.11.5301-5306.1998

Li, S., Prasanna, X., Salo, V.T., Vattulainen, I., Ikonen, E., 2019. An efficient auxin-inducible degron system with low basal degradation in human cells. Nat. Methods 16, 866–869. https://doi.org/10.1038/s41592-019-0512-x

Mendoza-Ochoa, G.I., Barrass, J.D., Terlouw, B.R., Maudlin, I.E., de Lucas, S., Sani, E., Aslanzadeh, V., Reid, J.A.E., Beggs, J.D., 2019. A fast and tuneable auxin-inducible degron for depletion of target proteins in budding yeast. Yeast Chichester Engl. 36, 75–81. https://doi.org/10.1002/yea.3362

Milholland, K.L., AbdelKhalek, A., Baker, K.M., Hoda, S., DeMarco, A.G., Naughton, N.H., Koeberlein, A.N., Lorenz, G.R., Anandasothy, K., Esperilla-Muñoz, A., Narayanan, S.K., Correa-Bordes, J., Briggs, S.D., Hall, M.C., 2023. Cdc14 phosphatase contributes to cell wall integrity and pathogenesis in Candida albicans. Front. Microbiol. 14.

Milholland, K.L., AbdelKhalek, A., Baker, K.M., Hoda, S., DeMarco, A.G., Naughton, N.H., Koeberlein, A.N., Lorenz, G.R., Anandasothy, K., Esperilla-Muñoz, A., Narayanan, S.K., Correa-Bordes, J., Briggs, S.D., Hall, M.C., 2022. Reduced Cdc14 phosphatase activity impairs septation, hyphal differentiation and pathogenesis and causes echinocandin hypersensitivity in Candida albicans. https://doi.org/10.1101/2022.09.29.510203

Mnaimneh, S., Davierwala, A.P., Haynes, J., Moffat, J., Peng, W.-T., Zhang, W., Yang, X., Pootoolal, J., Chua, G., Lopez, A., Trochesset, M., Morse, D., Krogan, N.J., Hiley, S.L., Li, Z., Morris, Q., Grigull, J., Mitsakakis, N., Roberts, C.J., Greenblatt, J.F., Boone, C., Kaiser, C.A., Andrews, B.J., Hughes, T.R., 2004. Exploration of Essential Gene Functions via Titratable Promoter Alleles. Cell 118, 31–44. https://doi.org/10.1016/j.cell.2004.06.013

Morawska, M., Ulrich, H.D., 2013. An expanded tool kit for the auxin-inducible degron system in budding yeast. Yeast Chichester Engl. 30, 341–51. https://doi.org/10.1002/yea.2967

Mulhern, S.M., Logue, M.E., Butler, G., 2006. Candida albicans Transcription Factor Ace2 Regulates Metabolism and Is Required for Filamentation in Hypoxic Conditions. Eukaryot. Cell 5, 2001–2013. https://doi.org/10.1128/ec.00155-06

Natsume, T., Kanemaki, M.T., 2017. Conditional Degrons for Controlling Protein Expression at the Protein Level. Annu. Rev. Genet. 51, 83–102. https://doi.org/10.1146/annurev-genet-120116-024656

Nicastro, R., Raucci, S., Michel, A.H., Stumpe, M., Garcia Osuna, G.M., Jaquenoud, M., Kornmann, B., De Virgilio, C., 2021. Indole-3-acetic acid is a physiological inhibitor of TORC1 in yeast. PLOS Genet. 17, e1009414. https://doi.org/10.1371/journal.pgen.1009414

Nishimura, K., Fukagawa, T., Takisawa, H., Kakimoto, T., Kanemaki, M., 2009. An auxin-based degron system for the rapid depletion of proteins in nonplant cells. Nat Meth 6, 917–922. http://www.nature.com/nmeth/journal/v6/n12/suppinfo/nmeth.1401_S1.html

Nishimura, K., Yamada, R., Hagihara, S., Iwasaki, R., Uchida, N., Kamura, T., Takahashi, K., Torii, K.U., Fukagawa, T., 2020. A super-sensitive auxin-inducible degron system with an engineered auxin-TIR1 pair. Nucleic Acids Res. 48, e108–e108. https://doi.org/10.1093/nar/gkaa748

Paul, S., McDonald, W.H., Moye-Rowley, W.S., 2018. Negative regulation of Candida glabrata Pdr1 by the deubiquitinase subunit Bre5 occurs in a ubiquitin independent manner. Mol. Microbiol. 110, 309–323. https://doi.org/10.1111/mmi.14109

Powers, B.L., Hall, M.C., 2017. Re-examining the role of Cdc14 phosphatase in reversal of Cdk phosphorylation during mitotic exit. J Cell Sci 130, 2673–2681. https://doi.org/10.1242/jcs.201012

Prozzillo, Y., Fattorini, G., Santopietro, M.V., Suglia, L., Ruggiero, A., Ferreri, D., Messina, G., 2020. Targeted Protein Degradation Tools: Overview and Future Perspectives. Biology 9. https://doi.org/10.3390/biology9120421

Prusty, R., Grisafi, P., Fink, G.R., 2004. The plant hormone indoleacetic acid induces invasive growth in Saccharomyces cerevisiae. Proc. Natl. Acad. Sci. 101, 4153–4157. https://doi.org/10.1073/pnas.0400659101

Reuß, O., Vik, Å., Kolter, R., Morschhäuser, J., 2004. The SAT1 flipper, an optimized tool for gene disruption in Candida albicans. Gene 341, 119–127. https://doi.org/10.1016/j.gene.2004.06.021

Roemer, T., Jiang, B., Davison, J., Ketela, T., Veillette, K., Breton, A., Tandia, F., Linteau, A., Sillaots, S., Marta, C., Martel, N., Veronneau, S., Lemieux, S., Kauffman, S., Becker, J., Storms, R., Boone, C., Bussey, H., 2003. Large-scale essential gene identification in Candida albicans and applications to antifungal drug discovery. Mol. Microbiol. 50, 167–181. https://doi.org/10.1046/j.1365-2958.2003.03697.x

Salehin, M., Bagchi, R., Estelle, M., 2015. SCFTIR1/AFB-Based Auxin Perception: Mechanism and Role in Plant Growth and Development. Plant Cell 27, 9–19. https://doi.org/10.1105/tpc.114.133744

Sathyan, K.M., McKenna, B.D., Anderson, W.D., Duarte, F.M., Core, L., Guertin, M.J., 2019. An improved auxin-inducible degron system preserves native protein levels and enables rapid and specific protein depletion. Genes Dev. 33, 1441–1455. https://doi.org/10.1101/gad.328237.119

Schreiber, S.L., 2021. The Rise of Molecular Glues. Cell 184, 3–9. https://doi.org/10.1016/j.cell.2020.12.020

Song, Y., Cheon, S.A., Lee, K.E., Lee, S.-Y., Lee, B.-K., Oh, D.-B., Kang, H.A., Kim, J.-Y., 2008. Role of the RAM Network in Cell Polarity and Hyphal Morphogenesis in Candida albicans. Mol. Biol. Cell 19, 5456–5477. https://doi.org/10.1091/mbc.e08-03-0272

Tanaka, S., Miyazawa-Onami, M., Iida, T., Araki, H., 2015. iAID: an improved auxin-inducible degron system for the construction of a ‘tight’ conditional mutant in the budding yeast Saccharomyces cerevisiae. Yeast 32, 567–581. https://doi.org/10.1002/yea.3080

Trost, M., Blattner, A.C., Lehner, C.F., 2016. Regulated protein depletion by the auxin-inducible degradation system in Drosophila melanogaster. Fly (Austin) 10, 35–46. https://doi.org/10.1080/19336934.2016.1168552

Uchida, N., Takahashi, K., Iwasaki, R., Yamada, R., Yoshimura, M., Endo, T.A., Kimura, S., Zhang, H., Nomoto, M., Tada, Y., Kinoshita, T., Itami, K., Hagihara, S., Torii, K.U., 2018. Chemical hijacking of auxin signaling with an engineered auxin–TIR1 pair. Nat. Chem. Biol. 14, 299–305. https://doi.org/10.1038/nchembio.2555

Watson, A.T., Hassell-Hart, S., Spencer, J., Carr, A.M., 2021. Rice (Oryza sativa) TIR1 and 5′adamantyl-IAA Significantly Improve the Auxin-Inducible Degron System in Schizosaccharomyces pombe. Genes 12. https://doi.org/10.3390/genes12060882

Wilson, B.R., Davis, D., Mitchell, A.P., 1999. Rapid Hypothesis Testing with Candida albicans through Gene Disruption with Short Homology Regions. J. Bacteriol. 181, 1868–1874. https://doi.org/10.1128/JB.181.6.1868-1874.1999

Yesbolatova, A., Natsume, T., Hayashi, K., Kanemaki, M.T., 2019. Generation of conditional auxin-inducible degron (AID) cells and tight control of degron-fused proteins using the degradation inhibitor auxinole. New Methods Extr. Funct. Mamm. Genome 164–165, 73–80. https://doi.org/10.1016/j.ymeth.2019.04.010

Yesbolatova, A., Saito, Y., Kitamoto, N., Makino-Itou, H., Ajima, R., Nakano, R., Nakaoka, H., Fukui, K., Gamo, K., Tominari, Y., Takeuchi, H., Saga, Y., Hayashi, K., Kanemaki, M.T., 2020. The auxin-inducible degron 2 technology provides sharp degradation control in yeast, mammalian cells, and mice. Nat. Commun. 11, 5701. https://doi.org/10.1038/s41467-020-19532-z

Yu, S., Paderu, P., Lee, A., Eirekat, S., Healey, K., Chen, L., Perlin, D.S., Zhao, Y., 2022. Histone Acetylation Regulator Gcn5 Mediates Drug Resistance and Virulence of Candida glabrata. Microbiol. Spectr. 10, e00963–22. https://doi:10.1128/spectrum.00963-22

Zhang, L., Ward, J.D., Cheng, Z., Dernburg, A.F., 2015. The auxin-inducible degradation (AID) system enables versatile conditional protein depletion in C. elegans. Development 142, 4374– 4384. https://doi.org/10.1242/dev.129635

